# Macrophage-associated lipin-1 transcriptional co-regulatory activity is involved in atherosclerosis

**DOI:** 10.1101/2020.06.02.130096

**Authors:** Cassidy M.R. Blackburn, Robert M. Schilke, Aimee E. Vozenilek, Brian N. Finck, Matthew D. Woolard

## Abstract

During atherosclerosis, macrophages engulf and break down deposited modified low-density lipoproteins (modLDLs) into lipids and free fatty acids. The lipids and free fatty acids from these modLDLs either need to be stored during a process called glycerolipid synthesis or broken down during β-oxidation. In addition, free fatty acids can activate transcription factors to promote a pro-resolving macrophage phenotype. The protein lipin-1 is involved in both glycerolipid synthesis and β-oxidation. Lipin-1 enzymatic activity is a key step in the glycerolipid synthesis pathway; lipin-1 transcriptional co-regulatory activity either augments or represses various transcription factors that are activated via free fatty acids that promote β-oxidation and inhibit inflammation. Lipin-1 enzymatic activity increases pro-inflammatory macrophage phenotypes and is atherogenic. In contrast, we have also demonstrated that lipin-1 transcriptional co-regulatory activity promotes pro-resolving macrophage phenotypes leading us to the hypothesis that lipin-1 transcriptional co-regulatory activity is atheroprotective. Using a mouse model to delete lipin-1 in myeloid cells, we have demonstrated that loss of lipin-1 increases plaque size and pro-inflammatory gene expression. We have also shown mice lacking lipin-1 in myeloid cells have increased plaque collagen deposition and larger necrotic core formation. Combined, these data suggest that though lipin-1 enzymatic activity is atherogenic, lipin-1 transcriptional co-regulatory activity is atheroprotective. Overall, the results suggest that the dual activities of lipin-1 contribute to atherosclerosis progression in opposite ways.

## Introduction

Atherosclerosis is a complex immunometabolic and chronic inflammatory disorder initiated by hypercholesterolemia [1-6]. Hypercholesterolemia promotes lipid deposition (in the form of low-density lipoprotein (LDL)) into the arterial intima. Within the arterial intima, LDLs become modified and cause immune cell recruitment and activation [7, 8]. In addition to cholesterol, LDLs contain lipids and free fatty acids. Macrophages initially attempt to clear excess modified low-density lipoproteins (modLDLs) and break down lipids and free fatty acids. Excess free fatty acids and lipids are incorporated into the glycerolipid synthetic pathway to be stored in lipid droplets along with cholesterol in the form of triacylglycerols (TAG), and phosphatidylcholines (PC) [1, 9, 10]. Free fatty acids are also broken down by beta-oxidation in the mitochondria [11]. The biosynthetic pathways that process free fatty acids drive macrophage immunological function. We have demonstrated that glycerolipid synthesis promotes macrophage pro-inflammatory responses and accelerates atherosclerosis [1]. However, free fatty acids bind to and activate numerous transcription factors including PPARs and LXRs that promote pro-resolving macrophage function and beta-oxidation within macrophages that can lead to atheroprotective responses.

Lipin-1 is a phosphatidic acid phosphohydrolase that converts phosphatidic acid into diacyglycerol during glycerolipid synthesis [1, 12-17]. Lipin-1 is also a transcriptional co-regulator that enhances the expression of genes related to beta-oxidation [18-21]. Lipin-1 binds and increase the activity of transcription factors PPAR-α, PGC-1α, PPAR-γ. Lipin-1 also binds but represses transcription factors NFAT and SREBP1 [19, 20, 22]. In macrophages, lipin-1 enzymatic activity promotes pro-inflammatory responses [1, 14, 16]. We have demonstrated that expression of a hypomorphic lipin-1 protein that lacks lipin-1 enzymatic activity within myeloid cells reduces atherosclerosis in an AAV8-PCSK9 model, suggesting that lipin-1 enzymatic activity is atherogenic [1]. In contrast, mice with a global deletion of lipin-1 have an increased atherosclerotic burden in a cholate model of disease, providing evidence that lipin-1 is atheroprotective [23]. The difference in atherosclerotic responses in these two-mouse models may be due to the contribution of lipin-1 within different tissues. However, there is the possibility that the difference in these responses is due to the different activities of lipin-1. We have previously demonstrated that lipin-1 can locate to the nucleus of macrophages with an atherosclerotic plaque, suggesting that lipin-1 transcriptional co-regulatory activity contributes to macrophage responses during atherosclerosis [14]. Nuclear lipin-1 contributes to IL-4 mediated macrophage responses and aids in wound healing [24]. Lipin-1 transcriptional co-regulatory activity augments PPARs to enhance anti-inflammatory responses and fatty acid oxidation genes in cells such hepatocytes and adipocytes [18, 20, 25, 26]. Free fatty acid breakdown promotes the activation of PPARs, which are transcription factors responsible for promoting macrophage wound-healing phenotypes to protect the lesion from rupturing [11, 20, 25-28]. Therefore, macrophage-associated lipin-1 transcriptional co-regulatory activity may be involved in atherosclerosis. We hypothesize that within modLDL-stimulated macrophages, lipin-1 transcriptional co-regulatory activity reduces atherosclerosis severity.

To evaluate the contribution of myeloid-associated lipin-1 transcriptional co-regulatory activity in atherosclerosis, we developed a mouse model with a myeloid-specific deletion of lipin-1 and induced atherosclerosis using the adeno-associated viral vector 8 expressing a gain-of-function mutation (D377Y) of mouse proprotein convertase subtilsin/kexin type 9 (referred to as AAV8-PCSK9) and high-fat diet[29]. Using this mouse model, we provide evidence that lipin-1 transcriptional co-regulatory in myeloid cells is atheroprotective.

## Methods

### Generation of bone marrow-derived macrophages (BMDMs)

BMDMs were generated as described previously [1]. Briefly, bone marrow cells were flushed from femurs of lipin-1 fl/fl (wild type) and lipin-1m KO mice with Dulbecco’s modified Eagle’s medium (DMEM; Gibco, 10829). Cells were centrifuged at 534xg for 5 minutes. Supernatant removed. Red blood cells in pellet were lysed with ammonium chloride-potassium carbonate lysis. After lysis, samples were filtered through 0.2μm cell strainer. Cells were resuspended in bone marrow differentiating media (KnockOut Dulbecco’s modified Eagle’s medium (DMEM; Gibco 10829) supplemented with 30% L-cell conditioned medium, 20% fetal bovine serum (Atlanta biologicals S11150), 2mM glutamax (Thermofisher 35050-061), 100U/ml penicillin-streptomycin (ATCC), 1mM sodium pyruvate (HyClone), and 0.2% sodium bicarbonate)and placed in non-treated T-75 flasks in incubator (5% CO2, 37 °C) until confluent. Once confluent (5-7days later), 11mM EDTA (pH=7.6) was added to flasks and incubated for 10 minutes on rocker. Cells were scraped off of flask, washed with 1x phosphate buffered saline (PBS), and centrifuged at 534xg to pellet and collect cells. BMDMs were resuspended in D-10 (DMEM (Gibco: 11965-092), 10% fetal bovine serum (Atlanta biologicals S11150), 2mM glutamax (Thermofischer 35050-061), 100U/mL penicillin/streptomycin (ATCC), and 1mM sodium pyruvate (HyClone) for further experiments

### L cell conditioned medium

Murine fibroblast cell line L929 (ATCC CCL-1) was grown in R-10 (RPMI 1640 (HyClone SH30027.01) supplemented with 10% FBS (Atlanta biologicals S11150), 1mM sodium pyruvate (HyClone) and 100U/ml penicillin/streptomycin)) 3.75×10^5^ cells were seeded in a T225 tissue culture treated flask with 75mLs of R-10 medium. Flasks were incubated at 37 °C, 5% CO_2_, for 12 days. After 12 days, medium was collected and centrifuged at 534xg 4 °C for 10 minutes to clear any cell debris. Supernatant was filtered (0.22mm), aliquoted, and stored at -80° C until use. As described previously [1]

### Subcellular fractionation

1.5×10^6^ BMDMs were seeded in non-treated dishes for 3 hours. A total of 6 million cells per treatment group was used. After 3 hours, media was removed and replaced with either D-10 (untreated) or D-10 + 50μg/ml oxidized LDL (hi oxLDL Kalen biomedical 770252-60) for 2 hours. After 2 hours, media was aspirated, cells were washed with PBS (no calcium or magnesium) and 750 uL of 11mM EDTA (pH=7.6) was added to cells. EDTA incubated on cells on rocker for 10 minutes. After 10 minutes, cells were scraped off and EDTA + cells were added to a tube. 1/3^rd^ of cells was added to two different 1.5ml microfuge tubes. One microfuge tube was spun at 9,600xg for 2 minutes to pellet cells. Supernatant was aspirated and cell pellet was placed in -80°C for at least 30 minutes (whole cell lysate fraction). The second microfuge tube was centrifuged 600xg for 5 minutes to gently pellet cells. Supernatant was gently aspirated and the last 1/3^rd^ of scraped cells was added to the tube. The tube was respun to pellet the rest of the cells and the supernatant was aspirated. The pellets were resuspended in lysis buffer: 1xRSB, 0.1M Sucrose buffer, 1% NP-40, 0.5% NaDOC+,1x protease inhibitor cocktail (Thermo Scientific), 1x phosphatase inhibitor cocktail 2 (Sigma Aldrich), and 1x phosphatase inhibitor cocktail 3 (Sigma Aldrich) and incubated for 15 minutes on ice. After incubation, microfuge tubes were vortexed 3 times for a few seconds each. Microfuge tubes were then centrifuged 900xg for 5 minutes at 4 °C. Supernatants were pipetted out and placed into clean microfuge tube labeled cytosolic fraction. Cytosolic fractions were lysed with equal volume of 2x NuPAGE lysis buffer (2X NuPage LDL sample buffer containing 100mM dithiothreitol (DTT; Life Technologies), 1x protease inhibitor cocktail (Thermo Scientific), 1x phosphatase inhibitor cocktail 2 (Sigma Aldrich), and 1x phosphatase inhibitor cocktail 3 (Sigma Aldrich) and stored at -20°C until Western blot. The translucent pellet (below the cytosolic fraction before it was removed) was washed with 1x RBS/0.1M buffer and repelleted via centrifugation. Supernatants were pipetted out and discarded. Remaining pellets are the nuclear fraction and were resuspended in 30uL RIPA (150mM Sodium Chloride, 1% NP-40, 0.5% sodium deoxycholate, 0.1% SDS, 50mM Tris, pH8) containing 1x protease inhibitor cocktail (Thermo Scientific), 1x phosphatase inhibitor cocktail 2 (Sigma Aldrich) and 1x phosphatase inhibitor cocktail 3 (Sigma Aldrich) and placed on ice for 20 minutes. The whole cell lysate fractions were removed out of the -80°C, thawed, and resuspended in 50uL RIPA buffer and placed on ice for 20 minutes. After 20 minutes, lysates were centrifuged at 16,000xg for 6.5 minutes to pellet unlysed particles (DNA, etc). Supernatants were removed and placed in a clean tube labeled either whole cell lysate or nuclear fraction and were lysed with equal volumes of 2x NuPAGE lysis buffer. Lysates were stored in -20°C until Western blotting

### Tissue processing for Western blot analysis

Tissues were harvested from mice and flash frozen in liquid nitrogen. Frozen tissues were stored in -80° C until use. 50mg tissue samples were lysed in 1mL RIPA buffer (150mM Sodium Chloride, 1% NP-40, 0.5% sodium deoxycholate, 0.1% SDS, 50mM Tris, pH8) containing 1x protease inhibitor cocktail (Thermo Scientific), 1x phosphatase inhibitor cocktail 2 (Sigma Aldrich) and 1x phosphatase inhibitor cocktail 3 (Sigma Aldrich) using tissue homogenizer from Bioject Inc. Homogenized tissues were incubated on ice for 30 minutes. Samples were then centrifuged at 16,000xg for 35 minutes at 4° C. Supernatant below the lipid layer but above the pellet was collected and transferred to clean 1.5ml microfuge tube and centrifuged at 16,000xg for 35 minutes at 4° C. Supernatant underneath the lipid layer was collected and transferred to a clean 1.5ml microfuge tube labeled tissue lysate. Tissue lysates were stored at -20° C until use.

### Western blot analysis

All tissue and cell lysate samples were sonicated and subsequently heat denatured at 95° C. Protein concentration of tissue lysate samples was determined by Pierce® 660 nm Protein Assay (Thermo Scientific). 40-50µg total protein of each tissue lysate sample and 20ug of cell lysate sample was added to 3.75µL4x NuPAGE buffer (106mM Tris HCl, 141mM Tris base, 2% SDS, 10% glycerol, 0.51mM EDTA, 0.22mM Brilliant Blue G250, 0.175mM Phenol Red), 0.75µL B-mercaptoethanol (Fisher Scientific) and water up to 15µL before loading. (Note: protein assay was not performed on the samples that underwent subcellular fractionation. After sonication and heat denaturation, 20uL of these samples were loaded into gel). Samples were placed in 95° C water bath for 7 minutes and then cooled to room temperature. Samples were centrifuged at 16,000xg for 1 minute before loading into gel. Samples were loaded into a 4 to 12% polyacrylamide NuPAGE Novex gel (Invitrogen). For Western blot in figure 1: 1x MOPs (50mM 4-morpholinepropanesulfonic acid, 50mM Tris base, 0.1% SDS, 1mM EDTA, pH7.7) running buffer was used to run gels. For Western blot in figure 2: 1xMES (20x MES: 50mM MES, 50mM Tris Base, 0.1% SDS, 1mM EDTA, water up to 1L) running buffer was used to run gels. Proteins were separated at 200 volts for 50 minutes. Semidry transfer was performed at 20 volts for 45 minutes onto a polyvinylidene difluoride (Immobilon-FL) membrane (EMD Millipore). Li-Cor Odyssey® blocking buffer (Li-Cor) was used to block membranes for 1 hour at room temperature. Primary antibodies were diluted 1:5000 (GADPH) or 1:1000 (Lipin-1) in 1% BSA in PBS with 0.01% sodium aside and incubated on membranes at 4° C overnight on a rocker. The membranes were then washed 4 times with 1x Tris-buffered saline and 0.1% Tween 20 (TBST) for 15 minutes each on a rocker after primary antibody incubation. Goat anti-Rabbit secondary antibody was diluted 1:2000 in 5% milk with TBST plus 0.01% SDS. Secondary antibodies were incubated on membranes for 2 hours at room temperature with rocking. The membranes were then washed 4 times with 1x TBST for 15 minutes each on a rocker. ImmunoCruz Western blotting luminol reagent (Santa Cruz sc-2048) was mixed and added to blots for 1 minute prior to exposure. Densitometry was performed using Image Quant TL Toolbox v8.2.0. Bands of interest were normalized to GAPDH for statistical analysis. Primary antibodies were as follows: Lipin-1 (Cell signaling technology 14906S) and GAPDH (Cell signaling technologies 2118S). Secondary antibodies were as follows: Rabbit Anti-Goat IgG (H+L) (Jackson ImmunoResearch 305-035-003) As previously described [29]

**Figure 1:**
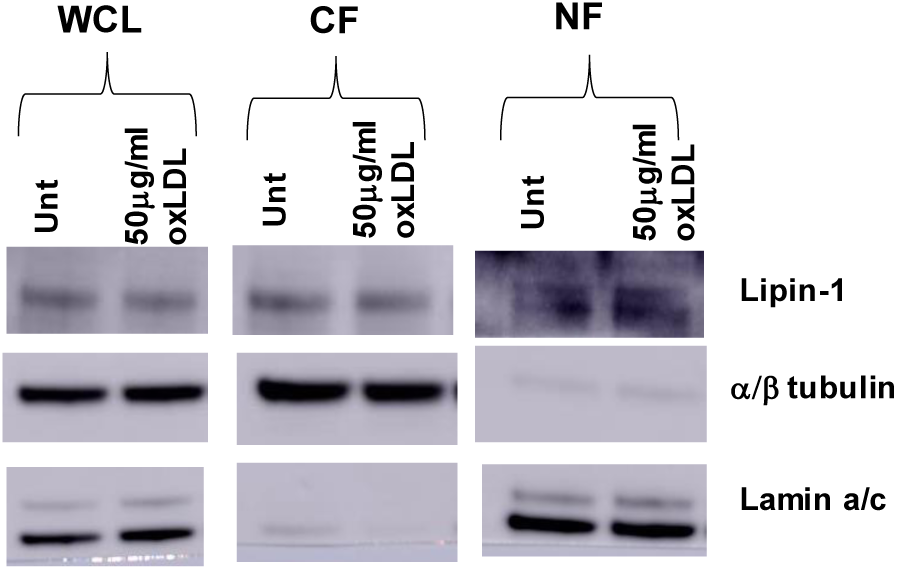
Lipin-1 localizes to the nucleus in bone marrow-derived macrophages. Femurs were isolated from wild type mice. Bone marrow was isolated from the femurs and differentiated into macrophages. Macrophages were seeded for 4 hours and then media was swapped with treatment for 2-4hours. Treatment was 50μg/ml oxLDL. D-10 media was used as untreated control. After treatment, cells were collected and split in half for whole cell lysate and the other half was fractionated into cytosolic and nuclear fractions. 20uL of protein lysate was loaded into Western blot and probed for lipin-1. α/β tubulin loaded as a cytosolic control and lamin a/c loaded as a nuclear control.

**Figure 2:**
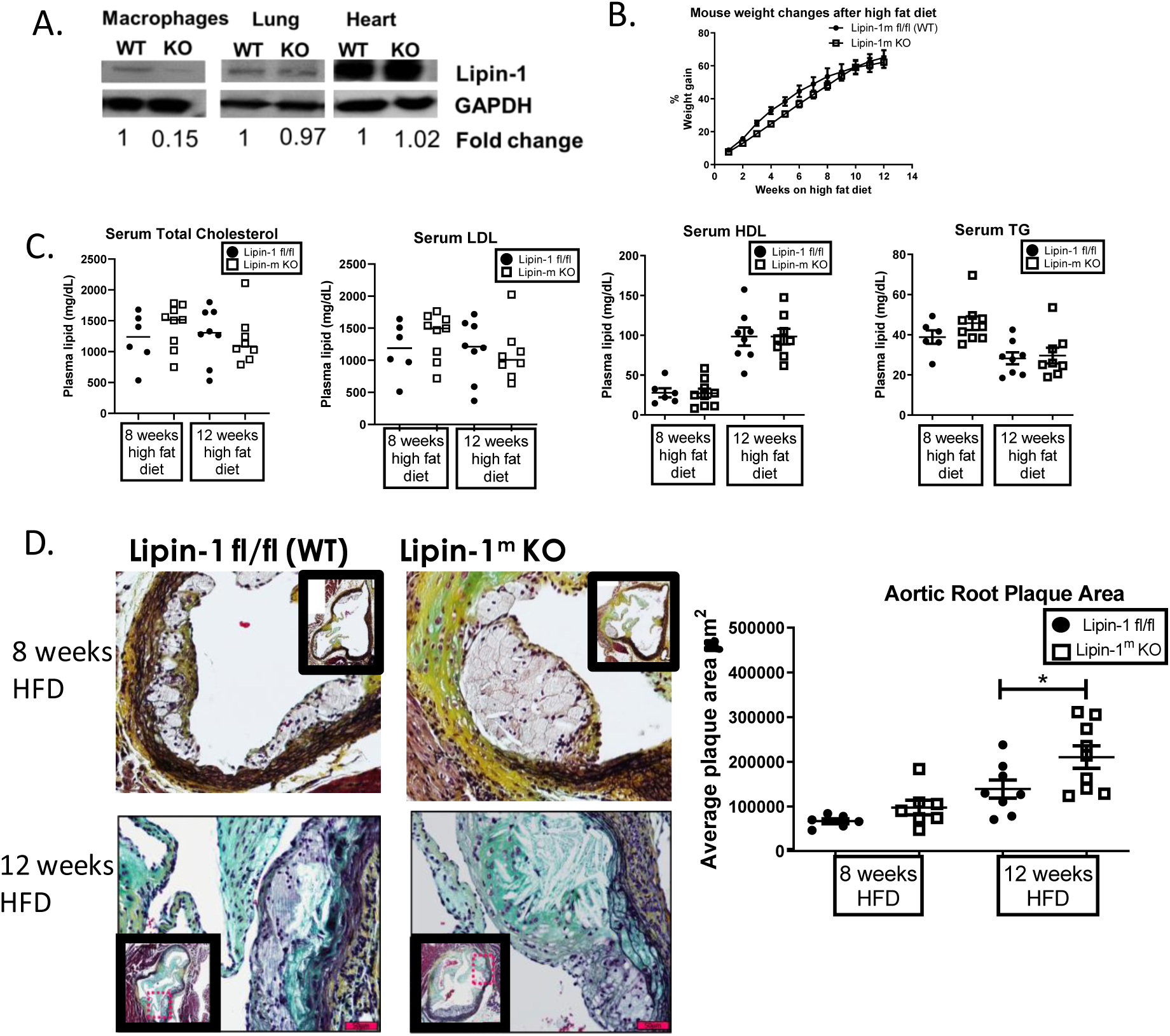
Lipin-1 transcriptional co-regulatory activity reduces atherosclerosis: A) Bone marrow-derived macrophages, lung, and heart tissues were collected and lysed from wild type (WT) and lipin-1m KO (KO) mice. Lysates were run on Western blot and probed for lipin-1 and GAPDH as a loading control. B-D: Wild type (Lipin-1 fl/fl) and myeloid-derived Lipin-1 knockout (Lipin-1^m^ KO) mice were injected with AAV8-PCSK9 to induce hypercholesterolemia and then fed high fat diet for 8, or 12 weeks. B) % weight gain over 8 or 12 weeks on high fat diet (HFD). C) Plasma lipid content after 8 or 12 weeks high fat diet. TC= total cholesterol; HDL= high density lipoprotein; TG= triglycerides; LDL= low-density lipoprotein. D) Aortic roots were harvested from mice, formalin fixed, paraffin embedded, sectioned and stained via MOVAT for plaque area analysis N=6-9 mice *p<0.5

### AAV8-PCSK9 viral vector preparation

Jacob Bentzon gifted DNA for the pAAV/D377Y-mPCSK9 (Addgene plasmid #58376 and it was packaged into adeno-associated virus serotype 8 (AAV8) using the helper and capsid plasmids from the University of Pennsylvania described in [1]. Viral stocks were sterilized via Millipore Millex-GV syringe filter (Billerica, MA). Post sterilization, viral stocks were titered via dot blot assay and then aliquoted and stored -80 ° C until use. Final product is referred to as AAV8-PCSK9.

### Animals and tissue harvest

Animal protocols were approved by LSU Health Sciences Center-Shreveport institutional animal care and use committee (protocol number P-19-029). All animals were cared for according to the National Institute of Health guidelines for the care and use of laboratory animals. Mice with a *Lpin1* allele with exon 7 of the *Lpin1* gene floxed with LoxP sites were generously provided by Brian Finck. To generate mice lacking lipin-1 in myeloid-derived cells only, the *Lpin1* floxed mice were crossed with LysM-Cre transgenic mice purchased from Jackson Laboratory (Bar Harbor, ME). Resulting offspring lacked lipin-1 within myeloid-derived cells (Lipin-1m KO mice). Lipin-1m KO mice were compared with the lipin-1m floxed littermate controls. Lipin-1mEnzy KO mice used are as described previously [1]. Briefly, mice with the *Lpin*1 gene floxed at exons 3 and 4 were crossed with LysM-Cre transgenic mice purchased from Jackson Laboratory to generate a mouse lacking myeloid-derived lipin-1 enzymatic activity but retaining the rest of the protein. 7-14 week old male mice were used for all studies. Male mice received retro-orbital injections of 3*10^10^ vector genomes of AAV8-PCSK9[29]. Immediately after AAV8-PCSK9 inoculation, mice were fed high fat, Western diet (TD 88137; Harlan-Teklad, Madison, WI) that contained 21% fat by weight (0.15% cholesterol and 19.5% casein without sodium cholate) for 8 or 12 weeks (duration of study). Mice were weighed once a week after beginning high fat diet. After 8 or 12 weeks, mice were anesthetized via isofluorane and subsequently euthanized via exsanguination and pneumothorax. Blood was collected via vena cava puncture and placed into heparinized blood collection tubes. Blood was then centrifuged at 5,500 rcf for 5 minutes; plasma was isolated and stored at -80°C until analysis. After blood isolation, mice were perfused with 1x phosphate buffered saline (PBS). Following perfusion, the heart, aorta, and carotids were isolated and stored in 10% formalin until analysis. As previously described [1]

### Blood analysis

Plasma from mice was analyzed for total cholesterol (Wako 999-02601), triglycerides (Pointe Scientific INC T7532-120), and high density lipoprotein (Wako 431-52501) using commercially available kits. Low density lipoproteins were calculated using the Friedwald equation. Any mice with total cholesterol or low density lipoprotein under 500mg/dL were not considered hypercholesterolemic and were eliminated from analysis. As previously described [1]

### Histology and Image Quantification

Hearts and aortic roots from mice were fixed in 10% neutral buffered formalin and embedded in paraffin. Aortic roots were sectioned into 5um sections. All sections were equally taken from anatomical landmarks of the initiation of inner leaflet valves. For plaque area analysis, aortic root tissue sections were stained via Russel-Movat pentachrome stain. Collagen composition within the plaque was determined using picrosirius red staining from established protocols. Stained tissues were imaged via Olympus DP72 camera and plaque area inside the internal elastic lamina was quantified and analyzed via CellSens software.

### Nanostring gene expression analysis

RNA was isolated from 12-48 aortic root tissue sections of 5μm each using the RNAstorm™ kit with the following modifications: tissue sections were scraped off of slides with a razor blade and placed in microfuge tubes, final elution was 15-20uL. RNA was analyzed for concentration and quality using the Qbit and Agilent. A minimum of 400ng total RNA at a minimum concentration of 20ng/ul was required with a minimum basepair region of 300 or more. Isolated RNA was shipped to Canopy Biosciences 4340 Duncan Ave, Suite 220, St. Louis MO 63110 to be analyzed for the predesigned myeloid innate immunity panel plus a spike in of about 30 genes. The resultant data was further analyzed by multiple T Test with assumption of constant standard deviation followed by Bonferroni-sidak correction for multiple T-tests using Graph Pad Analysis 8.0.

### Serum cytokine analysis

The concentration of pro-inflammatory cytokines was analyzed using the BioLegend LEGENDplex Mouse Inflammation Panel (13 plex) with V bottom plate (catalog number: 740446) according to the manufacturer’s instructions. Data were analyzed using the LEGENDplex Data Analysis Software.

### Immunofluorescence

Hearts and aortic roots from mice were fixed in 10% neutral buffered formalin and embedded in paraffin. Aortic roots were sectioned into 5μm sections. All sections were equally taken from anatomical landmarks of the initiation of inner leaflet valves. Tissue sections were rehydrated via placement in the following for 5 minutes each: 3 separate containers of xylene. 2 containers of 100% ethanol, 2 containers of 95% ethanol, 2 containers of 70% ethanol, 1 container of 50% ethanol, and 2 containers of deionized water. Antigen retrieval was performed via 20 minute incubation in 10mM citrate buffer in a microwave. Sections were cooled and then placed in blocking buffer (10% horse serum in heat denatured PBS +1% Bovine serum albumin (BSA) for at least one hour at room temperature. After blocking, tissue sections were rinsed with 1x TBST three times and then primary antibodies were added. Primary antibodies were incubated at 4° C overnight. After primary antibody incubation, tissue sections were rinsed in 1x TBST three times. Then tissue sections were incubated in 1%BSA in PBS for 5 minutes and removed. Secondary antibody was added to tissue sections and incubated for 2 hours at room temperature. After incubation, tissue sections were rinsed with 1xPBS twice. Tissue sections were then incubated with DAPI at a 1:50,000 dilution for 15 minutes in the humidifying chamber. DAPI was removed and a drop of n-propyl gallate was added. Slides were coverslipped and imaged using the Olympus camera and images were analyzed via CellSense software. Primary antibodies include: Rat-anti-Mac2 antibody (1:100) (Accurate Chem. CL8942AP)), Rabbit-anti-smooth muscle heavy chain II (1:400) (abcam ab53219), and donkey-anti-smooth muscle actin-Cy3 (1:200) (Sigma Aldrich C6198)). Secondary antibodies include: Alexa Fluor® 647donkey anti-rabbit IgG (A31573) and Alexa Fluor® 488 donkey anti-rat IgG (A21208) purchased from Life Technologies

### Necrotic Core formation analysis

Aortic root tissue sections were immunofluorescently stained as described above. Necrotic cores were quantified as areas of negative space (no staining) within the plaques.

### Statistical analysis

GraphPad Prism 8.0 (La Jolla, CA) was used for statistical analyses student T Test analysis was used for comparison between two data sets. All other statistical significance was determined using a one-way ANOVA with a Dunnett’s post-test. All experiments were performed a minimum of two times.

## Results

### Lipin-1 localizes to the nucleus in bone marrow-derived macrophages (BMDMs)

To determine if lipin-1 could be found in the nucleus of BMDMS, we performed a subcellular fraction of BMDMs followed by Western blot analysis. We confirmed successful fractionation by staining for α/β tubulin (cytosolic) and lamin a/c (nuclear). Minimal α/β tubulin was detected in the nuclear fraction and slight lamin a/c was detected in the cytosolic fraction indicating high enrichment between the fractions. Subcellular fractionation of BMDMs showed lipin-1 could be detected in both the cytosol and nuclear fraction (Figure 1). Treatment with oxidized low-density lipoprotein (oxLDL) also resulted in cytosolic and nuclear lipin-1 in BMDMs. These data confirm lipin-1 translocates to the nucleus in BMDMs and suggests modLDL treatment does not prevent lipin-1 localization into the nucleus (Figure 1). Lipin-1 exists as three different isoforms due to splice variants. The isoforms are lipin-1β, lipin-1-α, and lipin-1γ. Lipin-1γ has only been detected in the brain; however, lipin-1α, and lipin-1β have been detected in other hepatocytes, adipocytes, and macrophages [12, 13, 16, 17, 20, 24, 30]. There are two bands in the cytosolic fraction (one faint band and one dark band) and there are two prominent bands in the nuclear fraction for lipin-1. We would suggest these are the separate isoforms of lipin-1 being detected or different phosphorylation events.

### Lipin-1 transcriptional co-regulatory activity is involved in atherosclerosis

Mice lacking lipin-1 enzymatic activity in myeloid cells (Lipin-1mEnzy KO) had reduced plaque size compared to wild type mice suggesting lipin-1 enzymatic activity is atherogenic [1]. In contrast, we have demonstrated that lipin-1 transcriptional co-regulatory activity likely promotes wound healing responses in macrophages [24]. Wound-healing macrophages reduce atherosclerosis severity. Lipin-1 is found in the nucleus of macrophages within atherosclerotic plaques suggesting the transcriptional co-regulatory activity of lipin-1 is involved in atherosclerosis [14]. In order to determine the contribution of lipin-1 transcriptional co-regulatory activity to atherosclerosis, we developed a mouse lacking all of lipin-1 in myeloid-derived cells. Mice completely lacking lipin-1 from myeloid cells were generated by crossing mice with exon 7 of the *Lpin1* gene flanked by loxP sites with LysM-Cre transgenic mice. Deletion of exon 7 leads to a frameshift and premature stop codon with a complete loss of lipin-1 protein. Unfortunately, at this time, we are unable to generate a mouse model lacking only lipin-1 transcriptional co-regulatory activity because deleting this activity also diminishes enzymatic activity [20]. However, we are able to compare and contrast results with the previously published data with the lipin-1mEnzy KO mice to understand how lipin-1 transcriptional co-regulatory activity affects atherosclerosis. To confirm myeloid-specific deletion of lipin-1, protein was isolated from bone marrow-derived macrophages, lung tissue, and heart tissue from lipin-1m KO mice and wild type littermate controls. An 85% reduction of lipin-1 protein was observed in lipin-1m KO macrophages but equivalent levels of lipin-1 protein in the lung and heart tissue lysates was observed between wild type and lipin-1m KO mice (Figure 2A).

Having generated a mouse lacking lipin-1 from myeloid cells we investigated the contribution of myeloid-associated lipin-1 towards atherosclerosis using the AAV8-PCSK9 mouse model of atherosclerosis. Male lipin-1m KO mice and littermate control mice were injected with 3*10^10^ AAV8-PCSK9 viral genomes and then fed a high fat diet for 8 or 12 weeks to induce hypercholesterolemia and an atherosclerosis phenotype [29]. Lipin-1m fl/fl (wild type) and lipin-1m KO mice equally gained weight throughout the duration of the study (Figure 2B). Wild type and lipin-1m KO mice had equivalent total cholesterol, low density lipoproteins, high density lipoproteins and triglycerides in their serum indicating equal hypercholesterolemia (Figure 2C). Equal weight gain and serum lipids suggests both strains of mice were at equal risk of developing atherosclerosis; therefore, any differences in plaque size and composition would likely be due to the presence or absence of lipin-1. We examine plaque size of the aortic root as a marker of atherosclerotic severity in lipin-1mKO mice and littermate controls. Lipin-1m KO mice had larger plaques than wild type mice (Figure 2D). There was a slight increase in aortic root plaque area in lipin-1m KO mice after 8 weeks high fat diet and a significant increase in aortic root plaque area in lipin-1m KO mice after 12 weeks high fat diet as compared to littermate controls. These results are opposite of the results found in the lipin-1mEnzy KO mice. Lipin-1mEnzy KO mice had smaller plaques than wild type mice after both 8- and 12-weeks high fat diet [1]. Therefore, the loss of lipin-1 enzymatic activity reduces plaque size; however, this study demonstrates loss of both lipin-1 activities causes an increase in plaque size. Though the enzymatic activity of lipin-1 is atherogenic [1], these data suggest lipin-1 transcriptional co-regulatory activity is anti-atherogenic.

### Increased expression of collagen and pro-inflammatory genes in plaques of lipin-1m KO mice

The plaque size in the lipin-1m KO mice was nearly twice as large as the littermate control mice. Lipin-1m KO macrophages had decreased pro-resolving, wound healing macrophage polarization [24]. We hypothesized that myeloid-associated lipin-1 transcriptional co-regulatory activity promotes anti-inflammatory responses within the atherosclerotic plaque. We decided to examine changes in plaque gene expression between lipin-1mKO mice and littermate controls. We isolated mRNA from similarly positioned lesions within aortic roots of lipin-1m KO and littermate control mice on high fat diet for 8 and 12 weeks. Following mRNA isolation, we used Nanostring technology to determine changes in gene expression. We observed a significant increase in both pro-inflammatory gene expression and collagen gene expression in plaques of lipin-1mKO mice at both 8 and 12 weeks high fat diet compared to littermate controls. Collagen 12a1 mRNA and IL-23 mRNA were both upregulated in plaques from lipin-1m KO mice after 8- and 12-weeks high fat diet (Figure 3 A and B). By 12 weeks, collagen 11a1, 10a1, and 1a2 were also upregulated, as were other pro-inflammatory chemokines such as CCL28 and CCL12 along with IL-23. Increased collagen mRNA in a larger plaque can indicate a compensation mechanism where a fibrous cap is formed along the growing plaque to prevent plaque rupture. IL-23 can induce apoptosis and necrosis in cells within an atherosclerotic plaque; both processes promote necrotic core formation and thickening of the fibrous cap in the plaque.

**Figure 3:**
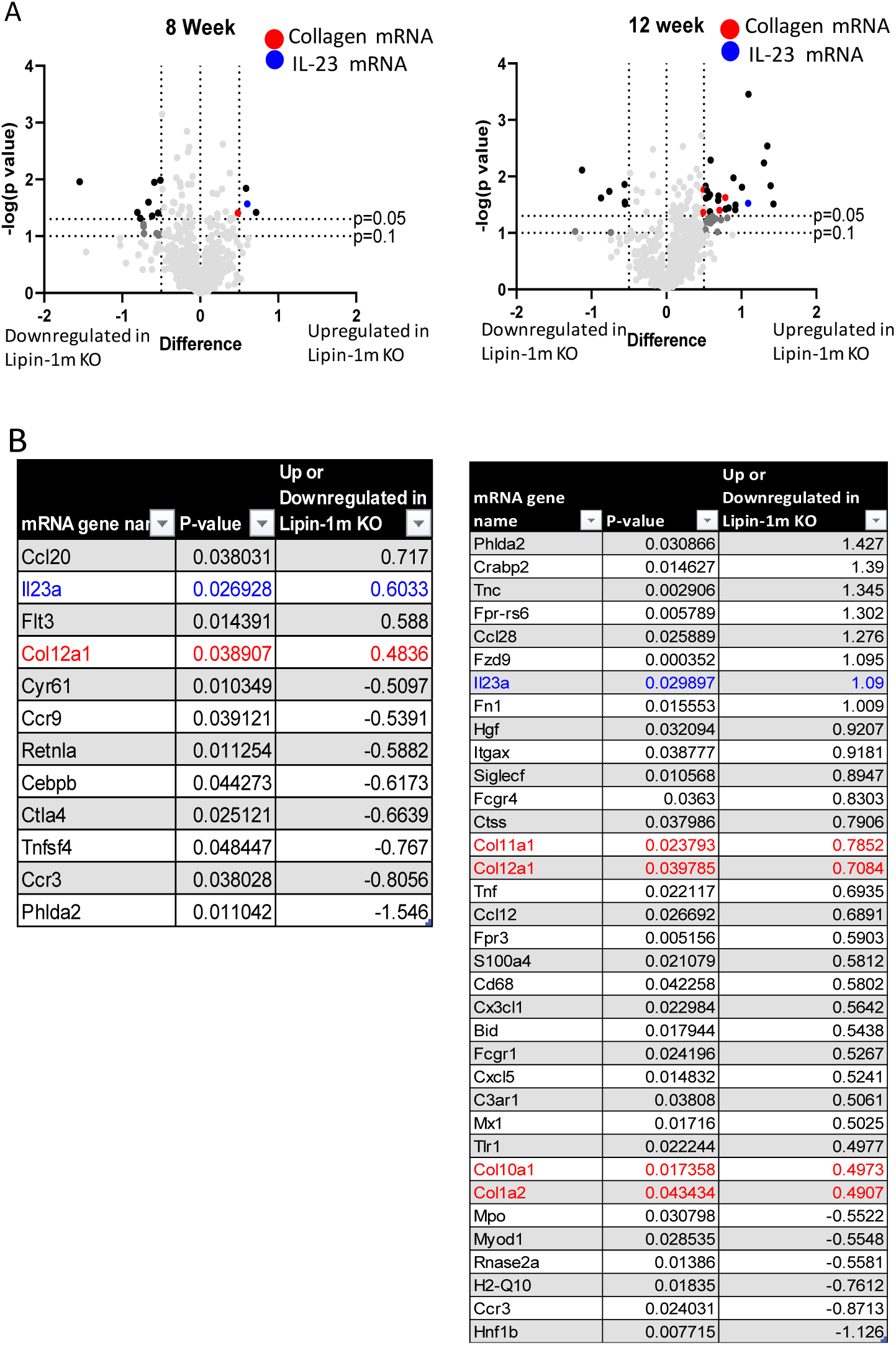
Lipin-1 regulates collagen and IL-23 gene expression. A-B) RNA was extracted from aortic root tissues sections from the wild type and lipin-1m KO mice after 8 or 12 weeks high fat diet described in figure 3. RNA was sent to Canopy biosciences for nanostring gene expression analysis. Up and downregulated genes are displayed for the lipin-1m KO mice compared to wild type mice. A) all genes plotted B) genes that are significantly up or downregulated 0.5 fold or more in the lipin-1m ko mice. n=6mice

Plaque gene expression data showed increased collagen mRNA in the lipin-1m KO plaques. Atherosclerotic plaque growth induces the collagen production to stabilize the plaque and prevent rupture [5, 6, 31, 32]. Increased collagen mRNA in the lipin-1m KO mice suggests increased collagen deposition. To quantify plaque collagen deposition, aortic root tissue sections from wild type and lipin-1m KO mice were stained with picrosirius red. Little to no collagen deposition was observed in plaques from mice on high fat diet for 8 weeks. By 12 weeks, however, lipin-1m KO mice had increased plaque collagen compared to wild type mice (Figure 4), supporting our mRNA gene expression data. Taken together, these data suggest that the loss of myeloid-associated lipin-1 accelerates plaque growth and corresponding collagen deposition to reduce potential plaque rupture.

**Figure 4:**
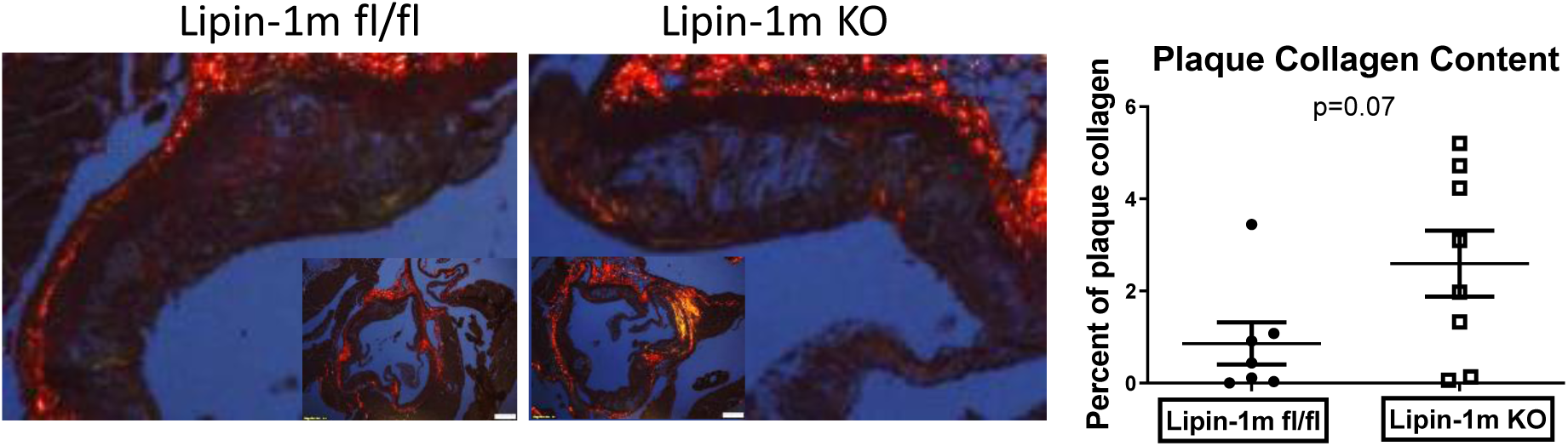
Myeloid-derived lipin-1 regulates plaque collagen deposition. Aortic root tissues sections from the wild type (Lipin-1 fl/fl) and lipin-1m KO mice after 12 weeks high fat diet described in figure 3 were stained with picrosirus red and analyzed for collagen deposition within the plaques. N=6-9 mice.

Gene expression data also showed increased IL-23 mRNA in the lipin-1m KO plaques after both 8- and 12-weeks high fat diet. Increased Il-23 mRNA should result in increased cytokine production that can be detected in the serum. Therefore, we quantified cytokine concentration in serum from the mice. We detected increased IL-23 cytokine concentration in the serum isolated from lipin-1m KO mice as compared to wild type mice at 12-week high fat diet (Figure 5). At 12 weeks high fat diet, we observed increased IFN-γ as well. When comparing lipin-1m KO and wild type mice regardless of time on high fat diet, we observed significant increases in IL-23, IL-27, IFN-γ and IL-1β. These cytokines are pro-inflammatory and pro-atherogenic. Thus, the loss of myeloid-derived lipin-1 results in increased plaque inflammation. The increase in plaque inflammation may be responsible for the increased collagen deposition in the lipin-1m KO plaques to increase fibrous cap thickness to prevent plaque rupture. Together, these results confirm our gene expression data and show increased plaque collagen deposition in the plaques with increased IL-23 and other pro-inflammatory cytokines in the mice. These data suggest more severe, inflammatory plaques in the lipin-1m KO mice.

**Figure 5:**
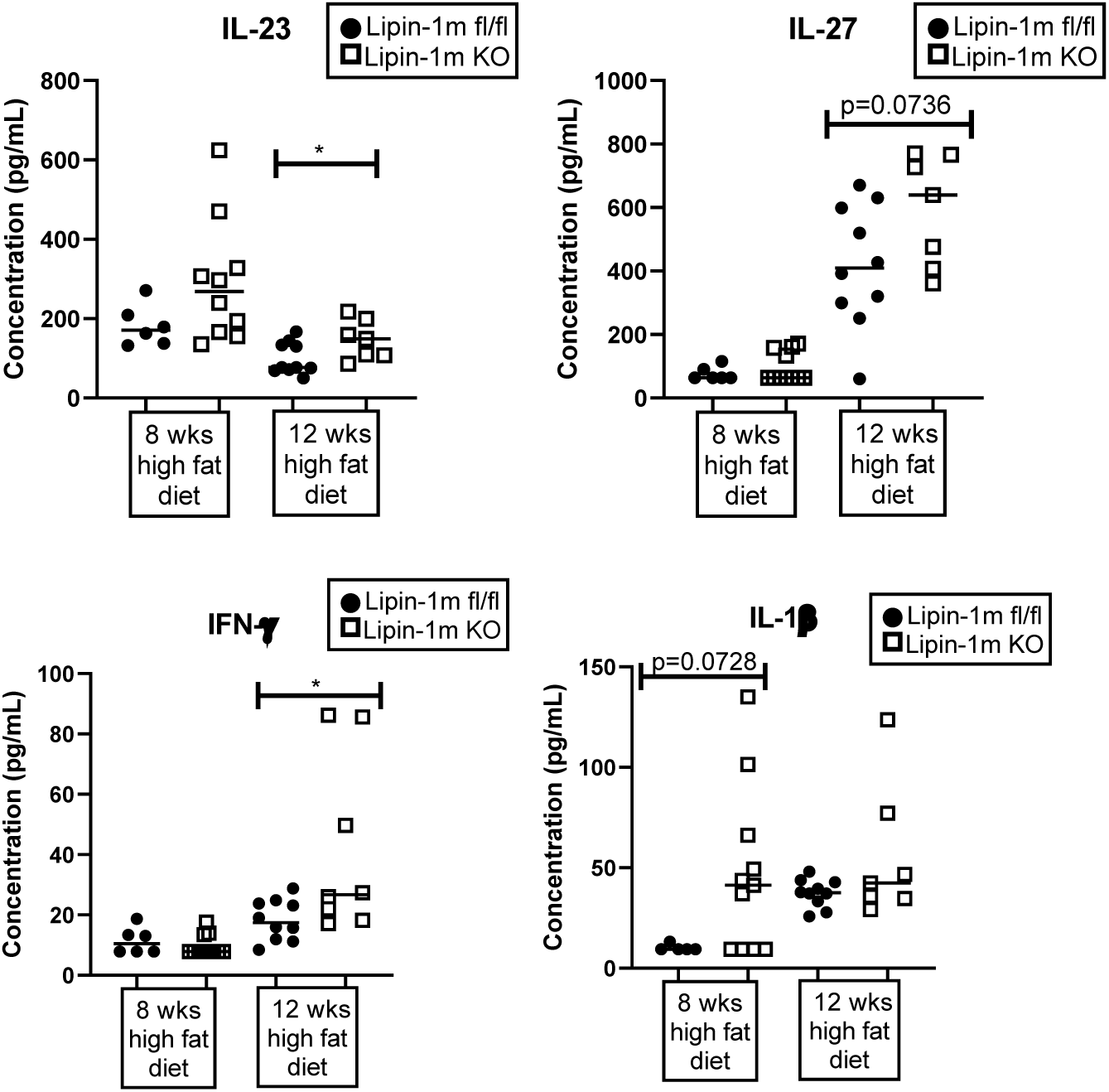
Myeloid-derived lipin-1 increases IL-23 and pro-inflammatory serum cytokines. After euthanasia, serum was collected from lipin-1m fl/fl (wild type) and lipin-1m KO mice on high fat diet for 8 or 12 weeks. Cytokines were measured via cytometric bead array. *= P<0.05 N=6-10 mice.

### Lipin-1 is involved in necrotic core formation

Vulnerable plaques are characterized by increased inflammation and increased apoptosis of interior macrophages and intimal smooth muscle cells[33]. Ineffective or incomplete apoptosis can cause cellular necrosis. Macrophages are responsible for clearing apoptotic cells and cellular debris from the result of necrosis. Macrophage-associated IL-23 promotes necrotic core formation within plaques [34]. We observed an increase in both IL-23 mRNA and serum IL-23. Thus far, our data have shown lipin-1m KO mice have larger plaques, increased collagen and IL-23 mRNA in the plaques, and increased IL-23 and other pro-inflammatory cytokines. These data led us to hypothesize that there would be larger necrotic cores in the plaques of lipin-1m KO mice. To determine necrotic core size, aortic root tissue sections from lipin-1m KO and littermate control mice were stained for macrophages, smooth muscle cells, and DAPI. We observed no difference in macrophage or smooth muscle cell content in the plaques between lipin-1m KO and littermate controls (Figure 6A). Necrotic cores were quantified as areas of negative space within the plaques. There was no difference in necrotic core formation between wild type and lipin-1m KO mice after 8 weeks high fat diet (Figure 6B). However, after 12 weeks of high fat diet, plaques in lipin-1m KO mice had larger necrotic cores than plaques in wild type mice (Figure 6B). Conversely, lipin-1mEnzy KO mice had no significant difference in necrotic cores compared to littermate control mice after 12 weeks high fat diet (Figure 6B). There was a significant increase in plaque necrotic core area of the lipin-1m KO (normalized to its wild type) compared to the lipin-1mEnzy KO (normalized to its wild type). Together, the data suggest that lipin-1 transcriptional co-regulatory activity regulates necrotic core formation, while lipin-1 enzymatic activity does not contribute to necrotic core formation.

**Figure 6:**
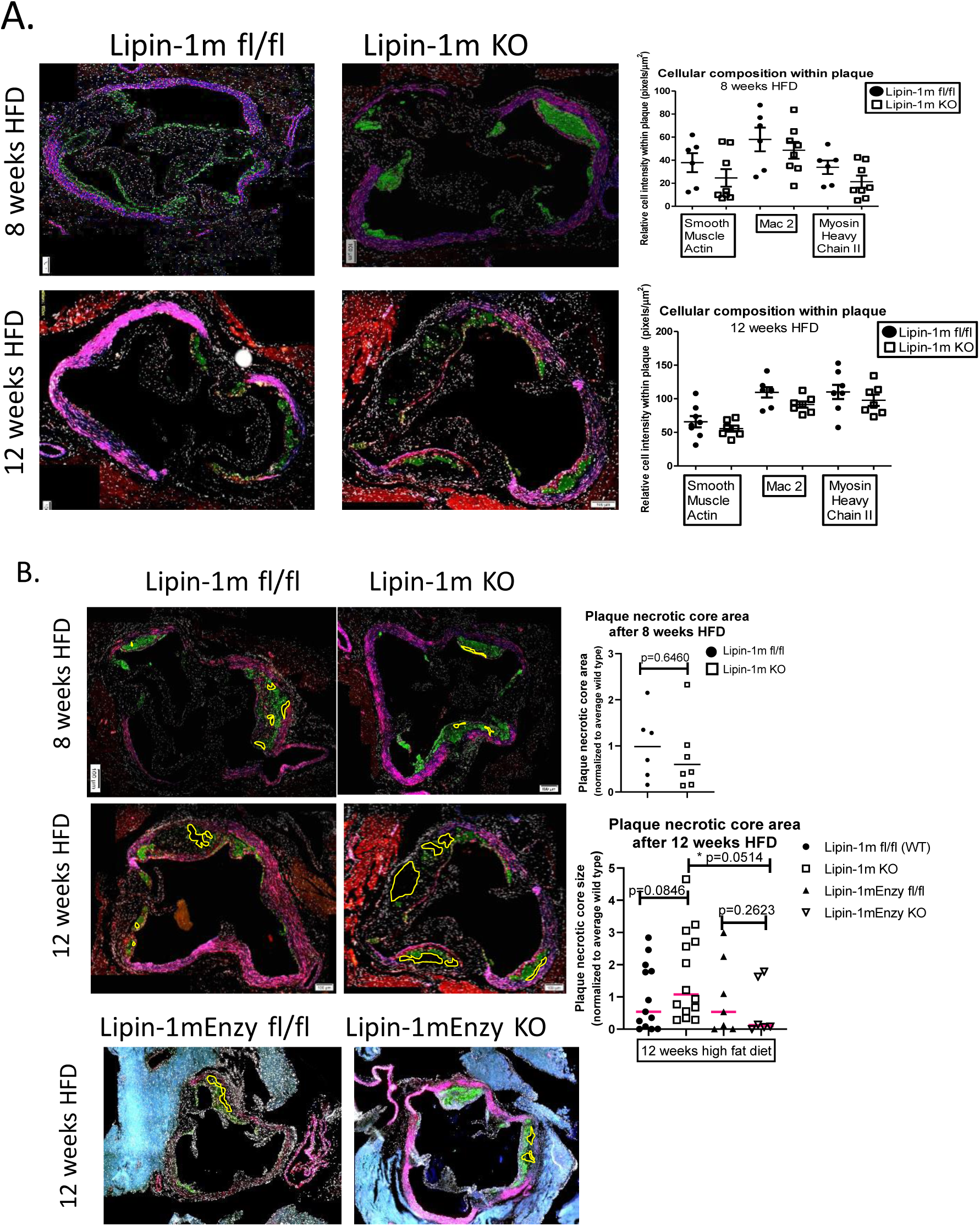
Lipin-1 transcriptional co-regulatory activity reduces necrotic core formation. **A)** Aortic root tissues sections from the wild type and lipin-1m KO mice after 8 or 12 weeks high fat diet described in figure 3 were immunoflurescently stained for macrophages (green) smooth muscle cells (blue and red) and DAPI (white). B) Mice described in A) plus wild type and lipin-1mEnzy KO mice induced with hypercholesterolemia via AAV8-PCSK9 and high fat diet for 12 weeks (described in [1]) were stained as described above. Necrotic core areas within plaques were quantified as areas with negative space (no staining) and are outlined in yellow. *p=<0.05 n=7-13 mice

## Discussion

Myeloid-derived lipin-1 phosphatidic acid phosphatase (enzymatic activity) is atherogenic. [1]. Lipin-1 has a separate, independent transcriptional co-regulatory activity that contributes to wound-healing macrophage polarization [1, 24]. These data suggested that lipin-1 transcriptional co-regulatory activity might be involved in atherosclerotic progression. In this study, we observed the loss of myeloid-derived lipin-1 results in increased atherosclerotic plaque size within the aortic root. Furthermore, we observed an increase in pro-inflammatory gene expression, collagen deposition, IL-23 and other pro-inflammatory cytokines, and necrotic core size within the lipin-1m KO mice. This phenotype contrasts our findings with the lipin-1mEnzy KO mice suggesting that each activity of lipin-1 contributes to atherosclerosis, but in opposite ways. These data suggest that lipin-1 transcriptional co-regulatory activity is atheroprotective.

The loss of myeloid associated-lipin-1 results in increased atherosclerosis severity; whereas the loss of myeloid associated-lipin-1 enzymatic activity results in decreased atherosclerosis severity. These data suggest that while lipin-1 enzymatic activity is atherogenic, lipin-1 transcriptional co-regulatory activity is atheroprotective. In hepatocytes and adipocytes, lipin-1 transcriptional co-regulatory activity increases the expression of anti-inflammatory and β-oxidation genes. In macrophages, anti-inflammatory and β-oxidation genes increase macrophage wound-healing responses, which may confer atheroprotection. Lipin-1 transcriptional co-regulatory activity also enhances skin wound healing; macrophage wound healing functions are important in reducing atherosclerosis severity [24]. In this study, we demonstrated that lipin-1m KO mice have increased plaque area and increased pro-inflammatory gene expression compared to wild type mice. These results are opposite of our previous work in which mice lacking enzymatic activity of lipin-1 had smaller plaques and reduced inflammation. Therefore, it is likely that the lack of lipin-1 transcriptional co-regulatory activity is responsible for the increase in plaque size and pro-inflammatory genes.

In the nucleus, lipin-1 binds and increases the activity of transcription factors. We have demonstrated that lipin-1 is found in the nucleus of unstimulated and modLDL-stimulated BMDMs. We also observed lipin-1 in the nucleus of plaque macrophages in humans [14]. Lipin-1 transcriptional co-regulatory activity augments the activity of the PPAR family of transcription factors. PPARs promote macrophage wound-healing responses which are atheroprotective [25, 26, 35]. PPAR-δ agonists have decreased atherosclerosis progression and reduced pro-inflammatory responses in macrophages [35]. Our data show that lipin-1m KO mice have increased plaque size with increased pro-inflammatory cytokines suggesting the lack of lipin-1 may result in reduced PPAR activity causing the increase in atherosclerosis severity. PPAR-γ agonists decrease IL-23 synthesis in mice after an allergen challenge [36]. While, PPAR-δ agonists decreased IL-23 in the central nervous system and lymphoid organs during multiple sclerosis and decreased renal necrosis under chronic ischemia [37, 38]. Lipin-1 also represses the activity of transcription factors such as NFAT4C and SREBP1 [19, 22]. These transcription factors promote pro-inflammatory macrophage responses. Combined, these studies suggest that potential for lipin-1 transcriptional co-regulatory activity promoting PPARs and inhibiting SREBP1 and NFAT4C may be an important mechanism during atherosclerosis progression.

In different inflammation models, PPAR agonists decrease IL-23 cytokine levels [37]. We see an increase in IL-23 mRNA within the plaque and cytokines in the serum of lipin-1m KO mice. IL-23 receptor on macrophages increases antigen-presenting capabilities in an autocrine manner, allowing the cytokine into lesion areas and increasing secretion of pro-inflammatory cytokines leading to pro-inflammatory macrophage polarization [39]. Human patients with carotid atherosclerosis had increased IL-23 in serum and increased serum IL-23 was correlated with increased disease progression and mortality [40]. The lipin-1m KO mice have increased disease progression demonstrated by larger plaques and increased pro-inflammatory cytokines such as IL-1β and IFN-γ. Furthermore, IL-23 production in mice increases macrophage apoptosis and necrosis [34]. Humans with advanced atherosclerosis have increased apoptotic and necrotic macrophages and smooth muscle cells in high rupture risk areas of plaques [33, 41, 42]. PPAR-δ is induced when macrophages engulf apoptotic cells and functions as a transcriptional sensor for dying cells to help increase apoptotic cell engulfment [43]. If programed apoptosis is unable to complete, cells then undergo necrosis. Increased necrosis leads to an increase in necrotic core formation within plaques. Necrotic core formation increases plaque inflammation thus potentiating atherosclerosis severity. Our data show lipin-1m KO mice have increased plaque IL-23 and larger necrotic cores than wild type or lipin-1mEnzy KO mice. These data suggest lipin-1 transcriptional co-regulatory activity may control or regulate necrotic core formation while lipin-1 enzymatic activity does not contribute to necrotic core formation. We propose that in modified LDL stimulated macrophages, lipin-1 translocates into the nucleus, binds transcription factors such as PPARs, resulting in decreased IL-23 secretion with reduced plaque growth and necrotic core formation thus reducing overall atherosclerosis severity (Model).

Overall, this study demonstrates macrophage-associated lipin-1 transcriptional co-regulatory activity contributes to reducing plaque growth, plaque inflammation, and necrotic core formation. The data obtained from this study could lead to enhanced therapeutics to specifically target plaque-induced inflammation during atherosclerosis. More work will need to be done to understand the exact mechanisms for how lipin-1 transcriptional co-regulatory activity regulates IL-23 production during atherosclerosis.

### Hypothetical Model

Macrophage engulfs modified LDL and breaks it down into free cholesterol and fatty acids. Upon activation and dephosphorylation, lipin-1 translocates to the nucleus where it binds transcription factors. Lipin-1 transcription factor binding results in decreased IL-23 synthesis under atherosclerotic conditions. Decreased IL-23 reduces necrotic core formation and atherosclerosis severity

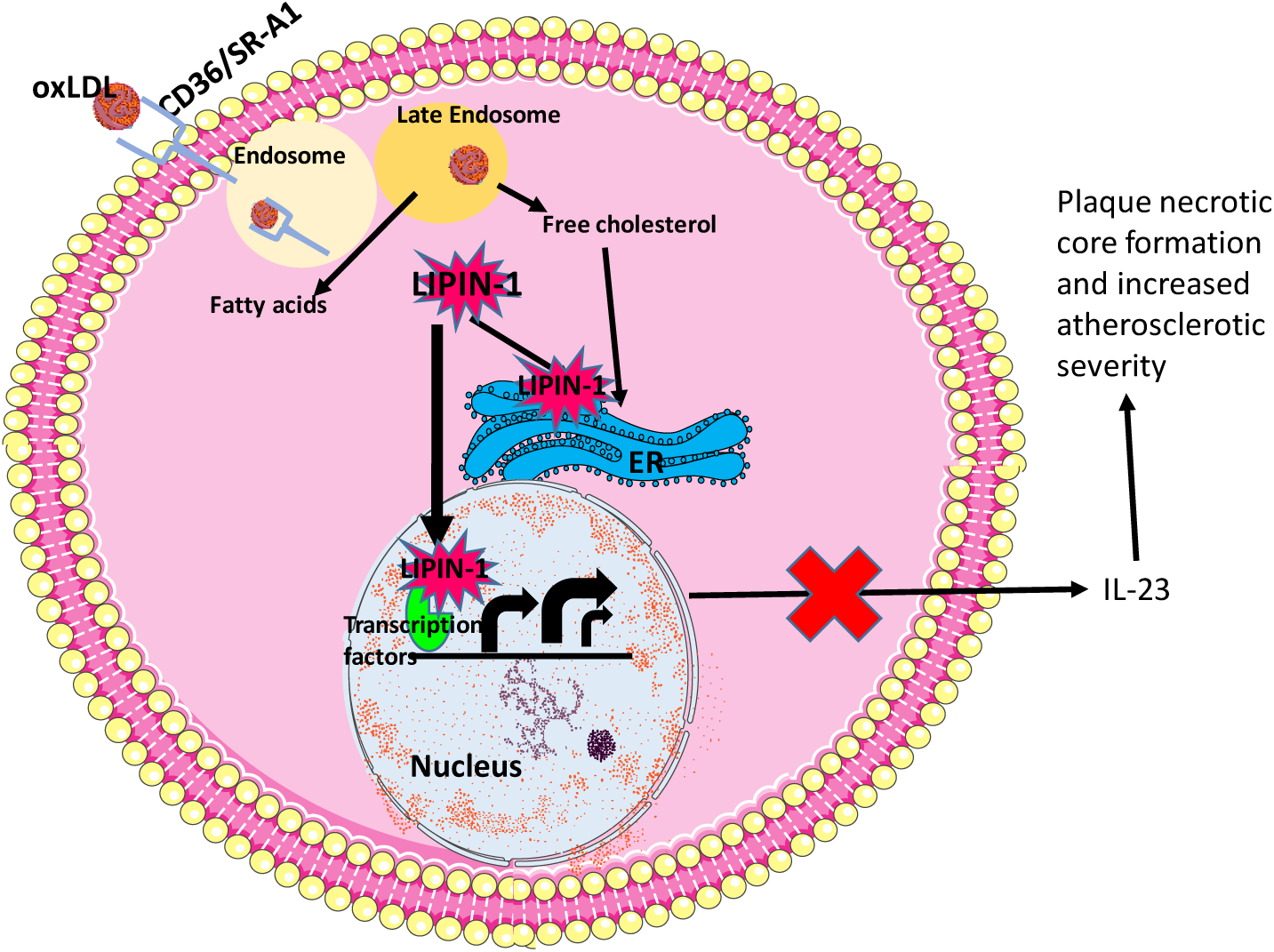

## Acknowledgements

We would like to thank Gabrielle Gahn, Faith Saxon, and Tyler Williams for determining genotypes of mice, Deshawn Blankenship for sectioning tissues used in this study, David Custis for his help in running serum samples for cytokine concentrations via flow cytometry, and Camille Abshire for helping verify sample concentrations and purity for the gene expression experiments.

